# Hypoxia modulates P-glycoprotein (P-gp) and breast cancer resistance protein (BCRP) drug transporters in brain endothelial cells of the developing human blood-brain barrier

**DOI:** 10.1101/2023.05.24.540054

**Authors:** Hafsah Mughis, Phetcharawan Lye, Guinever E. Imperio, Enrrico Bloise, Stephen G. Matthews

## Abstract

P-glycoprotein (P-gp) and Breast Cancer Resistance Protein (BCRP) are two multidrug resistance (MDR) transporters expressed at the blood-brain barrier (BBB). They confer protection against entry of harmful molecules into the fetal brain. The fetus develops under relatively low oxygen concentrations; however, pregnancy disorders (including pre-eclampsia) may lead to even lower intrauterine oxygen levels. We investigated the effects of hypoxia on transporter expression and activity in human fetal brain endothelial cells (hfBECs) isolated in early and mid-gestation. Results indicate decreased BCRP protein and activity under hypoxia in early-gestation hfBECs. Mid-gestation hfBECs exhibited an increase in P-gp and BCRP activity following hypoxia. Results suggest a hypoxia-induced reduction in fetal brain protection in early-pregnancy, but a potential increase in transporter-mediated protection at the BBB during mid-gestation. This would modify accumulation of various key P-gp and BCRP physiological and pharmacological substrates in the developing fetal brain and may play a role in the pathogenesis of neurodevelopmental disorders commonly associated with *in utero* hypoxia.

## Introduction

The developing blood-brain barrier (BBB) represents the primary barrier regulating transfer of key physiological substrates in/out of the brain and protect the developing brain against drugs and toxins present in the fetal circulation. The BBB is surrounded externally by pericytes and astrocyte foot processes and is composed of brain endothelial cells (BECs) which form the specialized brain microvasculature ^1,2^. The BECs express various proteins including tight junctions, and important multidrug resistance (MDR) transporters, P-glycoprotein (P-gp, encoded by the *ABCB1* gene) and Breast Cancer Resistance Protein (BCRP encoded by *ABCG2*). These transporters are responsible for the clearance of specific substrates out of the brain ^3^ and prevent the entry of neurotoxins and drugs, as well as endogenous molecules such as cytokines, steroids, and hormones into the brain parenchyma, supporting brain development and providing brain protection ^2,4–6^.

P-gp and BCRP are developmentally regulated at the BBB, with BCRP and P-gp expression being detected within the cerebral microvessels at 5- and 8-weeks post-conception, respectively ^7^. Recent studies have demonstrated that these transporters are functionally active in isolated human fetal BECs (hfBECs) derived from early and mid-gestation, ensuring the protection of the fetal brain as early as the first trimester of pregnancy ^8^. Their protective role is essential because approximately 70% of pregnant women are prescribed drugs ^5^, many of which are P-gp and BCRP substrates ^9,10^. However, there is very limited information about the effects of pregnancy complications on these transporters in the developing BBB, especially those involving fetal hypoxia.

Oxygen levels in first trimester are lower than later in pregnancy and slightly changes throughout gestation. Overall, oxygen concentrations in the embryo range between 2 – 9%, in vivo ^11^, whereas in the developing placenta, it ranges from 2.5 -3.3% oxygen before 10 weeks of pregnancy and rises to approximately 8% at 14 weeks of pregnancy, signifying the start of second trimester ^12,13^. By the end of the third trimester, placental oxygen levels drop to approximately 6% due to rapid fetal growth, which increases oxygen demands from the placenta ^12^. In normal pregnancies, mean oxygen concentrations at the end of third trimester are about 3.7% in the umbilical vein, and 2.5% in the umbilical artery ^12^. In this connection, the fetus under normal circumstances is able regulate oxygen in a way that the oxygen supply exceeds its metabolic demands. However, instances of oxygen deprivation or fetal hypoxia can cause critical injury to vital organs, especially the developing brain ^14,15^. Hypoxic stress in pregnancy is common and can occur in various situations including pregnancies at high altitude, maternal smoking, heart failure, anemia, placental insufficiency, placenta accrete, placenta previa or preeclampsia (PE) ^16^. Hypoxia at the BBB can lead to disruption in the distribution of water and ions, oxidative stress, and leakage of blood proteins into the brain. Hypoxia also activates various epigenetic mechanisms in the fetal brain that can lead to neurodevelopmental disruptions which impact brain function in adult offspring ^17^. Therefore, it is critical to understand the effects that hypoxia has at the developing BBB.

Very few studies have examined the effects of hypoxia on P-gp and BCRP at the BBB, and most have been conducted in the adult BBB in rat and mouse models ^18–25^. No studies have reported how hypoxia affects P-gp and BCRP, and therefore fetal neuroprotection, in the human fetal BBB during development. A number of different *in vitro* models of the BBB have been established, using guinea pig BECs ^26–28^, rat BECs ^29–31^, mouse models ^3,32,33^ and human 3D BBB models ^34–36^. However, these models are somewhat limited in terms of their physiological relevance towards the biology of human fetal BECs. In addition, oxygen concentrations used in these *in vivo* and *in vitro* hypoxia experiments are in the range of 3% -10%, which may not represent significant hypoxia for the developing brain ^37^. It has been reported that optimum (physiological) oxygen concentration for normal development of mammalian embryos is around 5% oxygen ^38^, and in most tissues during development, hypoxic responses occur around 0.5% -1% of oxygen ^37^.

We, therefore, hypothesized that hypoxia will modulate P-gp and BCRP expression and function in primary hfBECs derived from fetal brain microvessels in both early and mid-gestation. In the present study, we investigated the acute and chronic effects of hypoxia on the expression and function of P-gp and BCRP in *vitro*, in primary hfBECs derived in early (∼12 weeks) and mid (∼18 weeks) gestation, to determine whether hypoxia modifies the protective barrier function provided by P-gp and BCRP in the developing central nervous system (CNS). To our knowledge, this is the first study to examine the effects of hypoxia on the expression and function of P-gp and BCRP in developing hfBECs.

## Methods

### Cell culture and reagents

hfBECs were isolated from early and mid-gestation fetal brains as described previously ^8,39^. Briefly, fetal brains were collected after elective termination in early gestation and mid-gestation. All tissues were collected by the Research Centre for Women’s and Infants’ Health BioBank program at the Sinai Health System. Written informed consent (Protocol #18-0057-E) was acquired in adherence to the policies of the Sinai Health System and the University of Toronto Research Ethics Boards (REB). The REBs do not permit reporting or collection of any identifying or clinical information from elective pregnancy terminations, therefore we were unable to report on any clinical information of the donors.

hfBECs were grown in a 37°C/5% CO_2_-incubator in EndoGROTM-MV Complete Culture Media Kit®, (SCME004, Millipore, Blvd, ON, Canada) supplemented with recombinant human epidermal growth factor (5 ng/mL), L-Glutamine (10 mM), hydrocortisone hemisuccinate (1.0 μg/mL), heparin sulfate (0.75 U/mL), ascorbic acid (50 μg/mL), 20% Fetal Bovine Serum, penicillin (100 IU/mL), streptomycin (100 IU/mL) (15140-122, Life Technologies), and 1% normocin antibiotic (ant-nr-2, Invivogen, San Diego, CA, USA) at 20% oxygen (5% CO_2_, 37 °C). Following isolation, hfBECs were collected and stored in liquid nitrogen. All procedures, treatments, and analyses were performed with the cells at passage 4, due to limited number of cells available in prior passages ^39^. We have previously shown no differences in expression of P-gp and BCRP between passage 1 and passage 4 ^8^.

### Exposure of Human Fetal Brain Endothelial Cells (hfBECs) to hypoxia

Primary hfBECs derived in early gestation (11.3-12.5 weeks; *N=*6) and mid-gestation (17.2-18.5 weeks; *N=*6) were plated in 96-well plates (8,000 cells/well) for activity studies, or in 6-well plates (25000 cells/cm^2^) for gene and protein analysis and were grown to confluence, cultured for 24-hours at 20% oxygen (5% CO_2_, 37 °C) in EndoGROTM-MV Complete Culture Media (as described above). 24-hours after seeding, cells were challenged with different oxygen tensions: 20% (hyperoxia; control group), 5% (physiological fetal hypoxia) or 1% (fetal hypoxia) oxygen. Hypoxia experiments were undertaken using Modular Incubator Chambers (MIC-101, Billups-Rothenberg, Inc. California, USA). Three chambers were used for 20%, 5% and 1% oxygen treatments, and in each experiment, the respective chamber was flushed with 5% oxygen (5% oxygen, 5% CO_2_ and N_2_ balance) and 1% oxygen (1% oxygen, 5% CO_2_ and N_2_ balance) (Linde Canada), whenever opened. Cell-free media was infused with 1% oxygen and 5% oxygen for 24-hours prior to the experiment to equilibrate the media. For functional assays, culture media was removed and replaced with Dulbecco’s Modified Eagle Medium (DMEM) (21063029, Thermo Fisher Scientific), supplemented with 10% charcoal-stripped FBS (CS-FBS) (Wisent, Saint-Jean-Baptiste,QC, CA) overnight, before the start of the experiment. At 0-hours, the plates were then transferred to modular incubator chambers, which were flushed with 1% and 5% oxygen (5 min). Once flushed, the chambers were clamped and placed in an incubator (37 °C). Cells in the 20% oxygen group placed in a chamber the incubator with the clamps open. To confirm brain endothelial cells’ response to hypoxia, *VEGF* mRNA expression (a marker of hypoxia) was analyzed ^40^. In addition, an optical oxygen sensor (OXYLogger, PreSens; Regensburg, Germany) was placed in the 1% oxygen chamber, and oxygen measurements were logged throughout the experiment, to ensure no leakage in the chamber. All assays and analyses were performed at 3 different time-points 6-, 24-, and 48-hours; to observe the acute- and longer-term effects of hypoxia on P-gp/*ABCB1* and BCRP/*ABCG2* expression and function in the developing hfBECs.

### P-gp, BCRP and Esterase activity assays

P*-*gp function was assessed as described previously ^8,39,41^. Briefly, hfBECs (*N=*6/group, each treatment/subject run in triplicate) were seeded and treated as described above. On the day of the assay, cells were washed twice with warm (37°C) Tyrode salts solution (Sigma, #T2145) supplemented with sodium bicarbonate (1g/L, Sigma, #S6014). Cells were incubated with P-gp substrate calcein-acetoxymethyl ester (Ca-AM, 177831, 10^−6^ M, Sigma; 37°C, 5% CO_2_, 1 hour). Once substrate was added, the chambers containing the plates were flushed with 5% and 1% oxygen for 5 minutes again and sealed. After incubation (1-hour), plates were placed on ice, and cells were washed twice with ice-cold Tyrode salts solution and lysed in 200μL cold 1% Triton X-100 (Sigma #X100) in HBSS. Accumulation of Ca-AM was measured using a Microplate Reader at excitation/emission wavelengths of 485/510 nm.

Esterase activity was assessed, as described previously ^8,39,41^. Ca-AM is a non-fluorescing P-gp substrate that is cleaved by endogenous esterases upon entering the cells, being converted to fluorescent Calcein (which is not transported by P-gp). In cells that express P-gp, Ca-AM is transported out of the cell before this conversion ^42^. To confirm hypoxia does not affect esterase activity, hfBECs derived in mid-gestation (*N*=6) were exposed to hypoxia (1% oxygen; 6-, 24-, 48-hours). Cells were then washed once with warm Tyrode salts solution and incubated with 10^−6^ calcein-AM in 200uL warm Lysis buffer (1% Triton X-100 (Sigma #X100) in HBSS) for 1-hour under 1% oxygen. Conversion of calcein-AM to calcein, after treatment, was assessed using a Microplate Reader at excitation/emission wavelengths of 485/510 nm.

For BCRP, activity was assessed as described previously ^8,39,41^. Briefly, cells were incubated with 2μM chlorin e6 (Ce6, Santa Cruz Biotechnology, #SC-263067) for 1-hour at 37°C and 5% CO_2_. Ce6 is a photosensitizer that was identified as a specific BCRP substrate ^43,44^. In the absence of any inhibitors, Ce6 is effluxed by BCRP ^45^. After incubation (as described for P-gp above), plates were placed on ice, and cells were washed once with ice-cold Tyrode salts solution. Cells were lysed in 200μL cold 1% Triton X-100 (Sigma #X100) in HBSS. Accumulation of Ce6 was measured using a Microplate Reader at excitation/emission wavelengths of 407/667 nm.

To validate the specificity of substrates for their respective transporters, hfBECs were treated with inhibitors of P-gp and BCRP, verapamil (VPL) (100 μM; sigma, #V4629), and Ko143 (10 μM; sigma, #K2144), respectively ^8^. Inhibitor controls were included each time functional assays were performed.

### Protein analysis

Cells were harvested in lysis buffer (1 mol/L Tris-HCL pH 6.8, 2% SDS, and 10% glycerol) which included a protease and phosphatase inhibitor cocktail (78420, Thermo Scientific), and protein was extracted by sonication. Protein concentration was determined using the Pierce BCA Protein assay kit (Thermo Scientific). Total protein (22 μg) was loaded on 8% SDS polyacrylamide gels for electrophoretic separation (100V, 1.5h). Proteins were then transferred (10 min) from gels to polyvinylidene difluoride membranes (PVDF) using the Bio-Rad Trans-Blot® Turbo™ Transfer System. Membranes were blocked (1-hour, room temperature) with 5% skim milk in Tris-Buffered Saline containing 0.1% Tween (TBS-T), followed by incubation (4°C) with primary antibodies: P-gp (Abcam #ab170903, dilution 1:1000), and BCRP (Abcam #ab108312, dilution 1:1000, RRID: AB_10861951) for 40-hours; and the loading control ß-actin (Santa Cruz Biotechnology, #SC-47778, dilution 1:2000, RRID:AB_626632) for 12-hours. Membranes were subsequently washed (3x) with TBS-T and incubated (1-hour, room temperature) with HRP-linked anti-rabbit secondary antibody (1:10,000; GE Healthcare Bioscience, Baie d’Urfe, Canada). Chemiluminescence was assessed using the Luminata™ Crescendo Western HRP Substrate (Millipore) for all early-gestation samples. The SuperSignal™ West Femto (Thermo Scientific) Substrate was used for mid-gestation P-gp and BCRP samples. Membranes were probed with their respective substrates for 5 minutes and detected under UV using the Bio-Rad ChemiDoc™ MP Imaging system (RRID:SCR_019037). Protein bands were quantified by densitometric analysis using the Image Lab™ software (RRID:SCR_014210) and normalized against beta-actin signal for total protein assessment.

### Quantitative Real Time PCR (qPCR)

Total RNA was extracted using the RNeasy Mini Kit (73404, Qiagen, Toronto, Canada), following the manufacturer’s instructions. NanoDrop1000 Spectrophotometer (Thermo Scientific, Wilmington, USA) was used to determine the RNA purity/concentration (A260/A280 ratio of approximately 2.0), and total RNA (1 μg) was reverse transcribed to cDNA using iScriptTM Reverse Transcription Supermix (Bio-Rad, Mississauga, Canada). SYBR Green (Sigma-Aldrich) was used to run the quantitative polymerase chain reaction (qPCR) using the CFX 380 Real-Time system C1000 TM Thermal Cycler (Bio-Rad), with the following parameters: 1 cycle of 95°C for 2 mins, and 40 cycles of 95°C for 5s and 60°C for 20s. 5ng of RNA was used per reaction and each sample was set up in technical triplicates, where the cycle threshold (CT) value of each replicate per sample was recorded. A list of gene targets and primer sequences, with their references, is available in Table 1. The primers were purchased from Integrated DNA Technologies (Iowa, USA). The geometric mean of succinate dehydrogenase complex flavoprotein subunit A (*SDHA*) and cytochrome C1 (*CYC1*) was used to normalize the expression of target genes for all samples. The 2^-ΔΔCT^ Method ^46^ was used to calculate the relative mRNA expression of *VEGF, ABCB1* (encodes for P-gp), and *ABCG2* (encodes for BCRP).

**Table 1:**
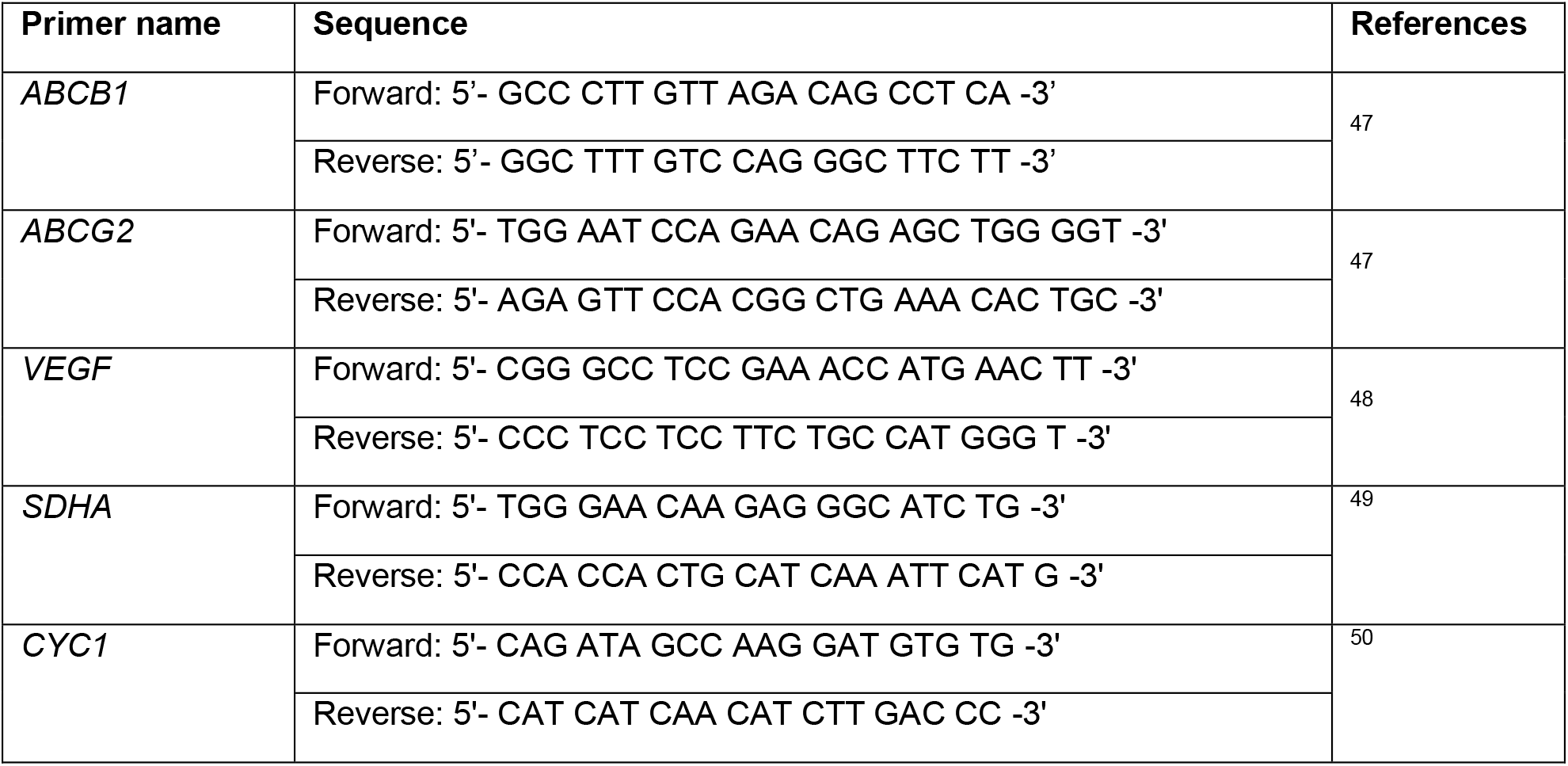
List of primers used in this study for qRT-PCR

### Cell Viability

Cell viability was measured using 6-well plates at 25000 cells/cm^2^ for each oxygen group (20%, 5% and 1% oxygen) at each time-point (6-, 24-, and 48-hours), as described above. After exposure, cells were washed with calcium-free HBSS, incubated with Accutase (1mL; 5 min., 37 °C, 5% CO_2_) and then mixed with cell media (2mL) to stop the reaction. An aliquot (100uL) of each well was mixed with equal parts trypan blue (100uL; 5 min) (15250-061; Gibco Life Technologies), mounted onto a hemocytometer (Hausser Scientific, Pennsylvania, United States) and cells were counted under the microscope. Cells which stained blue were deemed dead and those that extruded the dye, alive. Percent viable cells was determined using the following equation: % viable cells = viable cells/ (viable cells + dead cells) x 100 ^8^.

### Statistical Analyses

Statistical analyses were performed using GraphPad Prism (Inc., San Diego, USA) software version 9. Outliers were identified using the Grubbs’ test and normality was assessed using Shapiro-Wilk test. Gene and protein expression was analyzed using two-way ANOVA, followed by Dunnett’s multiple comparisons test, comparing between different oxygen tensions at each time-point in a single analysis. For P-gp and BCRP functional assays, a one-way ANOVA was used followed by Dunnett’s multiple comparisons test because data for each time-point was analyzed separately. Data are presented as mean ± standard error of the mean (S.E.M.). Differences were considered significant when P < 0.05.

## Results

### Hypoxia upregulates vascular endothelial growth factor (*VEGF)* in developing hfBECs

*VEGF* was used as a marker of the hypoxic environment in early and mid-gestation hfBECs to different levels of oxygen ^40^. *VEGF* mRNA levels were upregulated under 1% hypoxia at 24-(p<0.05) and 48h (p<0.01) in early gestation hfBECs (Figure 1A). In mid-gestation, *VEGF* mRNA levels in hfBECs were significantly upregulated under 1% oxygen, compared to 20% oxygen, at 6-hours (P<0.001), 24-hours (P<0.01), and 48-hours (P<0.05) (Figure 1B). Results from the oxygen sensor (Supplementary Figure 1A-B) confirm that oxygen concentrations were maintained throughout the treatment periods. The different oxygen tensions did not affect cell viability at any of the time points investigated in early gestation hfBECs (Supplementary Figure 1C-D).

**Figure 1:**
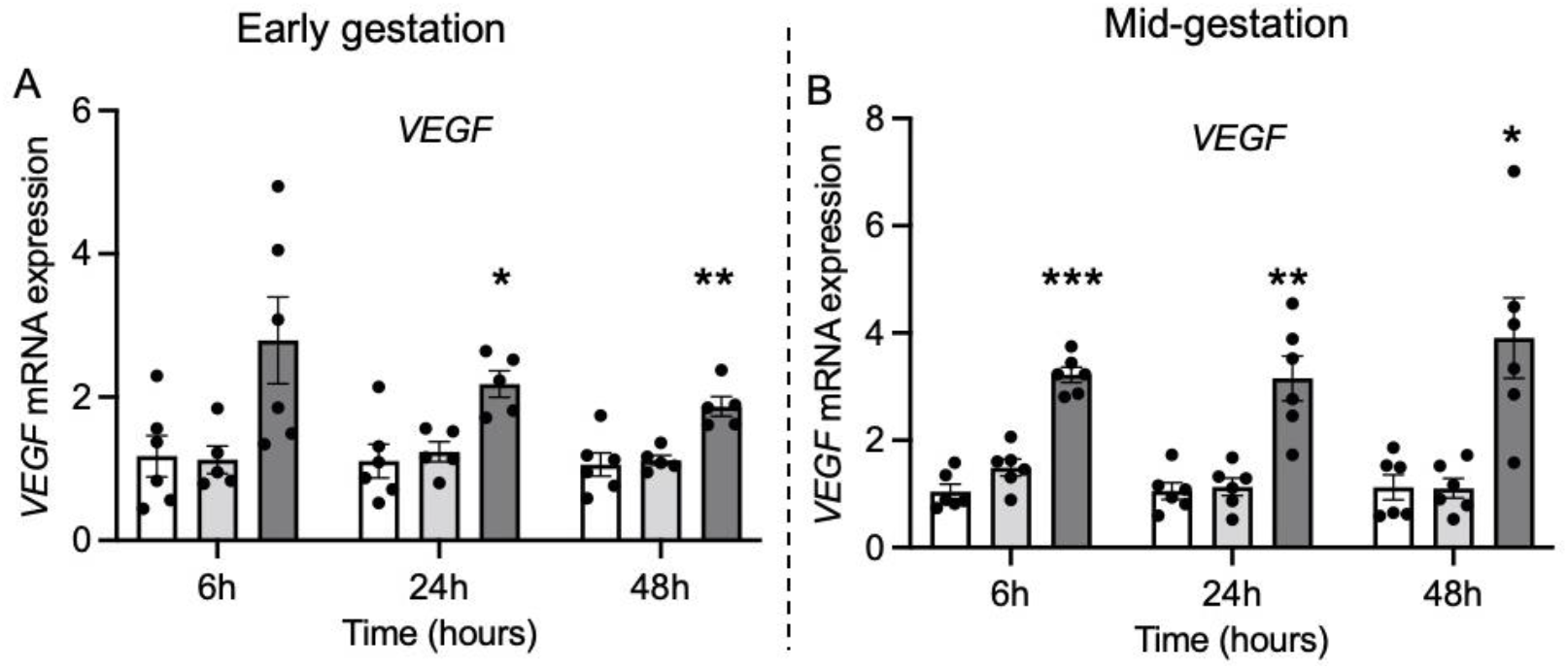
Hypoxia upregulates *VEGF* mRNA levels in human fetal brain endothelial cells (hfBECs) derived in early- and mid-gestation. Relative *VEGF* mRNA levels in early (A) and mid- (B) gestation hfBECs following 6-, 24-, and 48-hours of hypoxia (*N=*6/group). Statistical analysis: Two-way ANOVA followed by Dunnett’s multiple comparisons test. Values are displayed as mean ± S.E.M. Significant difference from 20% oxygen was set at (*) P<0.05; (**) P<0.01; and (***) P<0.001.

### Hypoxia promotes gestational-age-specific changes in Pgp and BCRP activity in hfBECs

In BECs derived in early gestation, there was no significant effect of 1% and 5% oxygen after 4, 24 and 48h on P-gp activity relative to 20% oxygen, however, there was a trend (P<0.08) towards an increase in P-gp activity under 1% oxygen at 48h (Figure 2A-C). In BECs derived in mid-gestation, there was a significant increase in P-gp activity after exposure to 1% oxygen compared to 20% oxygen at 6-hours (P<0.001, Figure 2D), 24-hours (P<0.05, Figure 2E), and 48-hours (P<0.01, Figure 2F). Interestingly, longer-term exposure (48h) to 5% oxygen led to a significant increase in P-gp activity (Figure 2F).

**Figure 2:**
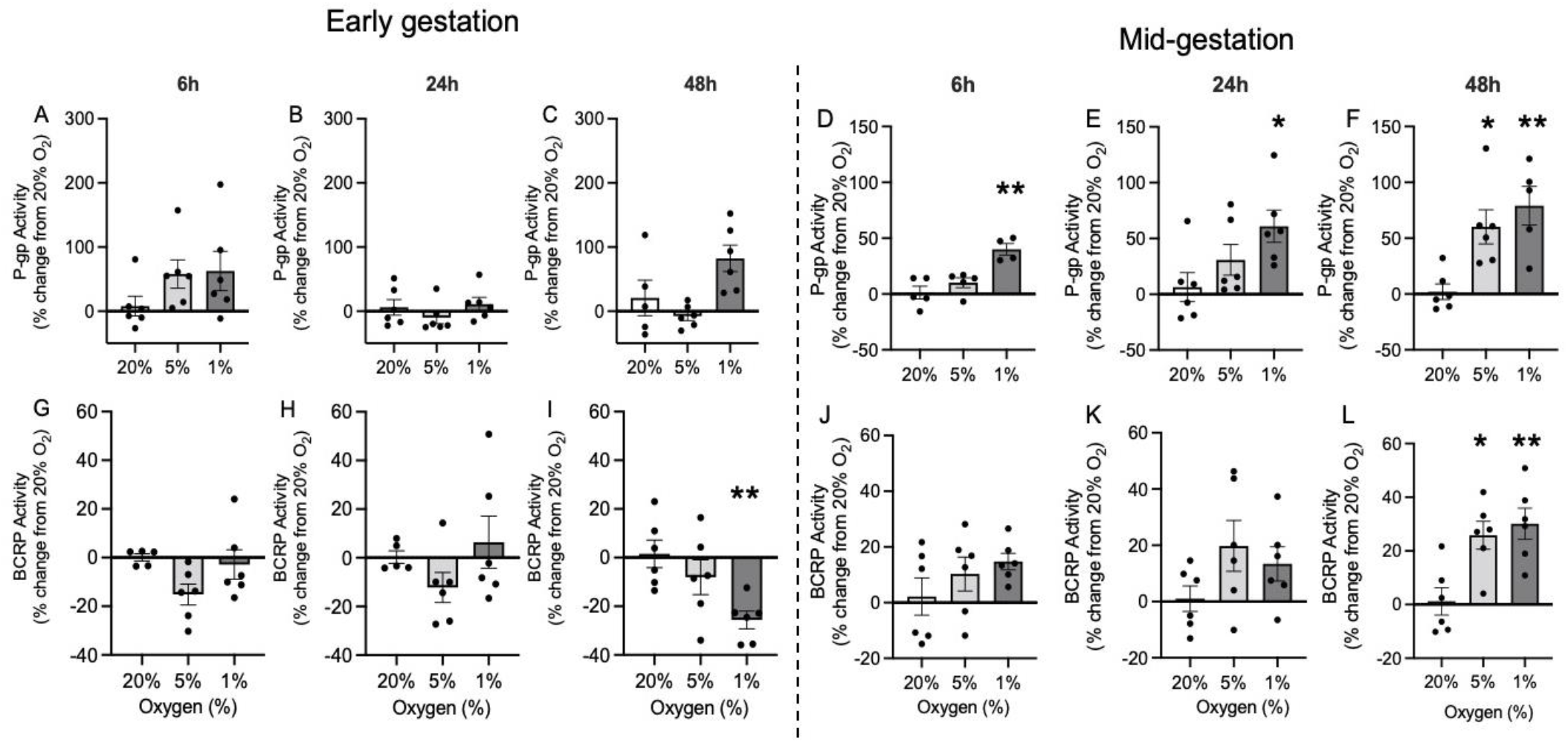
Hypoxia decreased BCRP activity in early gestation, and increased P-gp and BCRP activity in mid-gestation human fetal brain endothelial cells (hfBECs). Percent change in P-gp activity following hypoxia after **(A**,**D)** 6-hours, **(B**,**E)** 24-hours, and **(C**,**F)** 48-hours, and BCRP activity after **(G**,**J)** 6-hours, **(H**,**K)** 24-hours, and **(I**,**L)** 48-hours in early- and mid-gestation hfBECs (*N=*4-6/group, if *N*<6, an outlier has been removed). Activity is displayed as change relative to cells under 20% oxygen (control cells represented by a solid line at zero). Statistical analysis: One-way ANOVA followed by Dunnett’s multiple comparisons test. Values are displayed as mean ± S.E.M. Significant difference from 20% oxygen was set at (*) P<0.05; and (**) P<0.01.

BCRP activity was also affected by hypoxia in hfBECs. In early gestation, there was a significant decrease in BCRP activity after 1% oxygen exposure compared to 20% oxygen, at 48-hours (P<0.01, Figure 2I), whereas no differences were detected at other time points (Figure 2G & H). In contrast, in hfBECs derived in mid-gestation, there was a significant increase in BCRP activity after exposure to 1% oxygen (P<0.01) and 5% oxygen (P<0.05), compared to 20% oxygen, at 48-hours (Figure 2L). No significant differences were observed in BCRP activity after exposure to 1% and 5% oxygen at 6-hours and 24-hours (Figure 2J & K). Of importance, we also treated the developing hfBECs with specific P-gp (verapamil -VPL) and BCRP (KO-143) inhibitors. VPL increased calcein-AM accumulation, whereas Ko143 increased Ce6 accumulation in early and mid-gestation hfBECs in all oxygen treatments and time points investigated (Supplementary Figure 2A-L), validating the functional assays. Since the P-gp activity assay depends on the actions of cellular esterase ^41,42^, it was important to determine that hypoxia does not impact cellular esterase activity. We observed no changes in esterase activity following 6-, 24-, and 48-hours of hypoxia (1% oxygen) in mid-gestation hfBECs (Supplementary Figure 3).

### Hypoxia has specific effects modulating P-gp and BCRP protein and mRNA expression

We evaluated the effects of oxygen tension in modulating P-gp/*ABCB1* and BCRP/*ABCG2* expression in developing hfBECs. Hypoxia had no effect on P-gp protein levels at any of the time-points investigated in early and mid-gestation hfBECs (Figure 3A-D). *ABCB1* mRNA levels did not change in early hfBECs (Figure 4A) but were significantly decreased in mid-gestation hfBECs after exposure to 1% oxygen (P<0.05), compared to 20% oxygen at 24- and 48-hours (Figure 4B). BCRP protein levels were significantly decreased (P<0.01) after longer-term (48-hours) severe hypoxia (1% oxygen) compared to 20% oxygen in early-gestation hfBECs (Figure 3A & E) but did not change in hfBECs derived in mid-gestation (Figure 3B & F). No significant differences were observed in *ABCG2* mRNA levels under 5% or 1% oxygen at any time-point or gestational-age investigated (Figure 4C & D).

**Figure 3:**
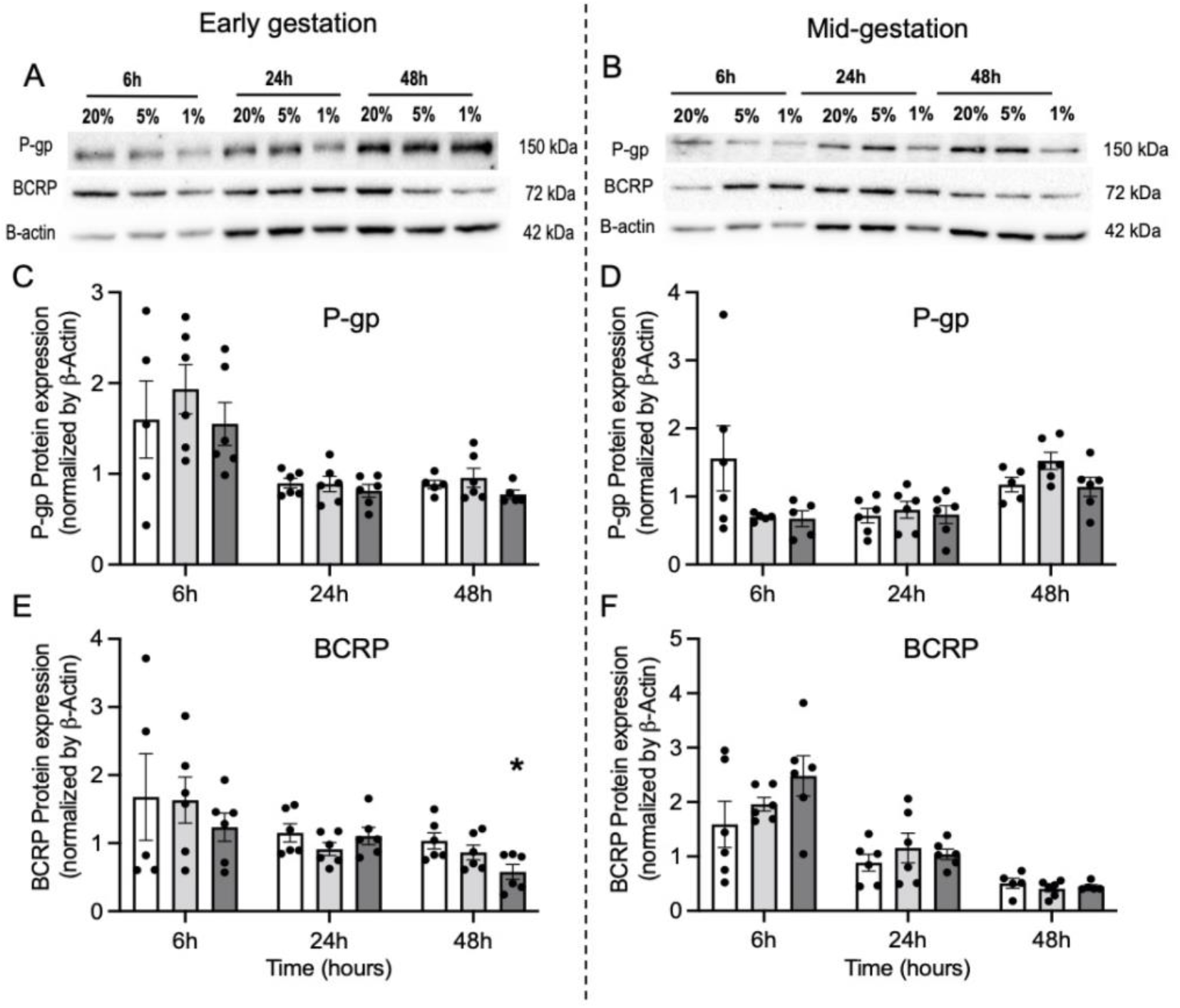
Hypoxia decreased BCRP protein levels in early gestation human fetal brain endothelial cells (hfBECs). (**A-B**) Representative P-gp and BCRP Western blot images, (**C-F**) densitometric analysis of P-gp and BCRP total protein levels, normalized to beta-actin (loading control for total protein), in early- and mid-gestation hfBECs following 6-, 24-, and 48-hours of hypoxia (*N=*5-6/group, if *N*<6, an outlier has been removed). Statistical analysis: Two-way ANOVA followed by Dunnett’s multiple comparisons test. Values are displayed as mean ± S.E.M. Significant difference from 20% oxygen was set at (*) P<0.05.

**Figure 4:**
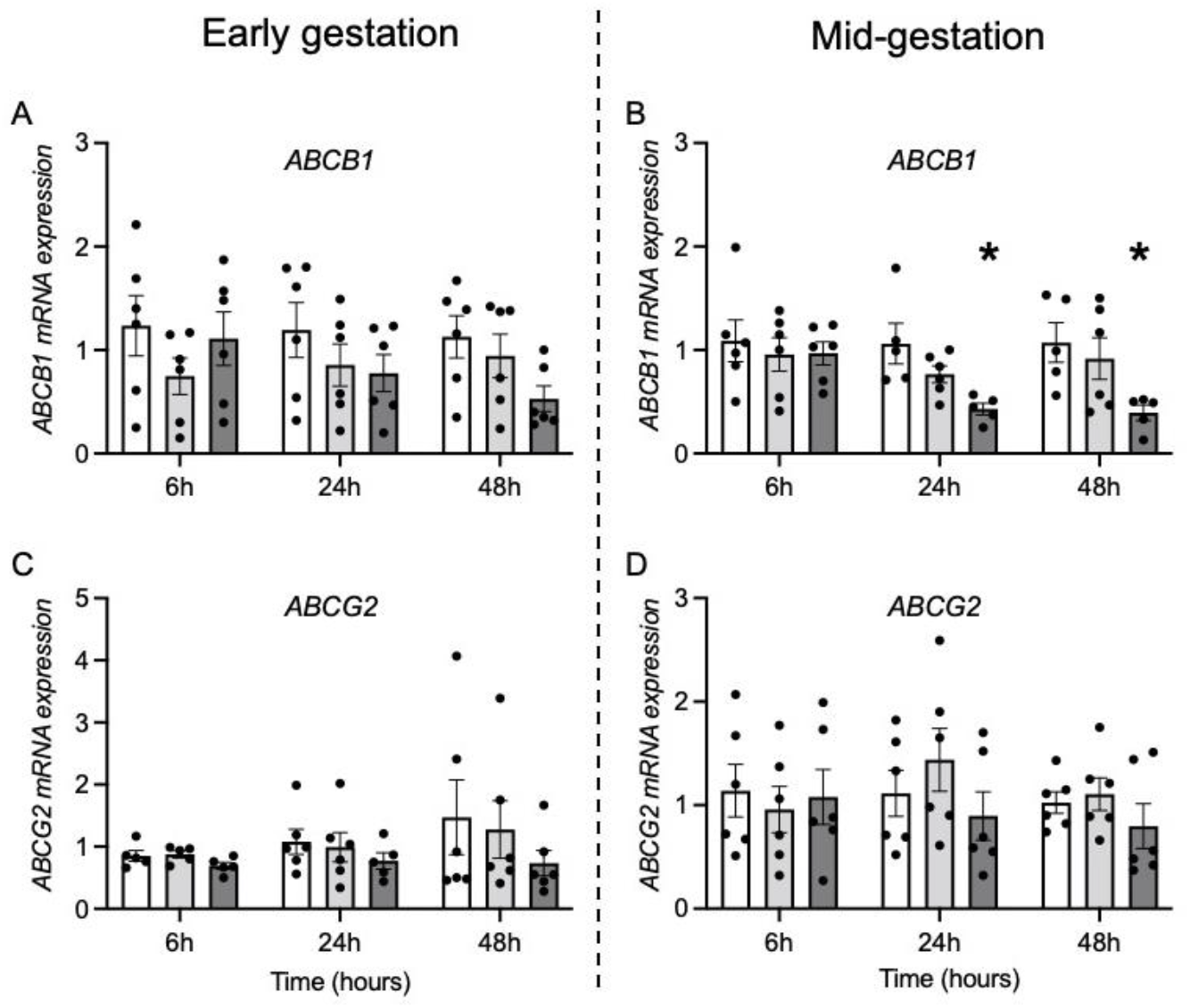
Effect of hypoxia on *ABCB1* and *ABCG2* mRNA levels in human fetal brain endothelial cells (hfBECs) derived in early- and mid-gestation. Relative *ABCB1* and *ABCG2* mRNA levels in early (A and C) and mid- (B and D) gestation hfBECs following 6-, 24-, and 48-hours of hypoxia (*N=*5-6/group, if *N*<6, an outlier has been removed). Statistical analysis: Two-way ANOVA followed by Dunnett’s multiple comparisons test. Values are displayed as mean ± S.E.M. Significant difference from 20% oxygen was set at (*) P<0.05.

## Discussion

In the present study, we have demonstrated for the first time, the effects of hypoxia on P-gp and BCRP expression and function in the developing human BBB using hfBECs derived from fetuses in early and mid-gestation as an experimental model. This follows careful characterization of the phenotype of the hfBECs used in this study. This involved determination of the expression and localization of important BEC markers such as Von Willebrand factor (vWF), zonula occludens (ZO)-1, and claudin (CLDN)5, as well as the functional expression of P-gp and BCRP, and the capability of hfBECs to form tube-like structures ^8^.

Research focused on the effects of hypoxia on P-gp and BCRP in BECs has given rise to contradictory results, depending on the disease model, the type of hypoxic treatment and the species ^18–22,24,25,29,51–53^. A study using bovine BECs showed downregulation of *ABCB1* mRNA after 4-hours of oxygen-glucose deprivation followed by 24-hours of reperfusion ^21^. Conversely, increases in P-gp protein expression were observed in an *in vitro* model of primary rat BECs following 6-hours of hypoxia via administration of hydrogen peroxide ^29^. Spudich et al. ^53^ subjected mice to middle cerebral artery occlusion and reported an increase in P-gp expression and activity. Using an *in vitro* model of the BBB (HBEC-5i -a cerebral microvascular endothelial cell line), BCRP protein expression decreased by 22% following sustained hypoxia (1% oxygen) ^24^, whereas in a rat *in vivo* stroke model (reversible occlusion of the right middle cerebral artery), there was a significant upregulation of *ABCG2* mRNA and BCRP protein levels 3-14 days after induction ^52^. In another *in vitro* study using adult human primary BECs, there was no effect of 1-, 4- or 8-hours of hypoxia (< 2% oxygen) on BCRP expression ^25^. To our knowledge, no studies have examined the effects of low oxygen tension, specifically on MDR transporter expression and activity in the human fetal BBB from early and mid-gestation.

There were no effects of hypoxia on the expression and function of P-gp in early gestation hfBECs, suggesting that fetal brain protection provided by P-gp at the developing BBB is likely maintained during hypoxic insults in early pregnancy. In contrast, hypoxia resulted in a significant and consistent decrease in BCRP protein expression and activity in hfBECs from early gestation, indicating that severe hypoxia may lead to an increase in the exposure to BCRP substrates in hypoxia-related pregnancy complications. These include toxins and other xenobiotics present in the maternal and fetal circulations that are harmful for the fetus, as well as endogenous factors such as folate and sphingolipids, that regulate neurodevelopment. The overall effects of hypoxia on transporter activity were more pronounced in mid-gestation hfBECs. Acute and longer-term hypoxia increased P-gp function. As such, it is possible that fetal hypoxia may result in an increase in P-gp-mediated efflux decreasing the transfer of various substrates across the BBB and conferring greater drug and toxin protection of the developing brain. However, at the same time, there would be a decrease in the brain accumulation of physiological substrates relevant to brain development (discussed below).

To understand how the increase in P-gp activity is regulated, P-gp protein and *ABCB1* mRNA levels were assessed under the same oxygen concentrations and time-points. P-gp protein showed no significant differences in expression at any time-point whereas *ABCB1* mRNA levels significantly decreased after 24- and 48-hours. This disconnect has been reported previously ^8,41,54–56^. Imperio et al. ^41^ examined the effect of infection mimics on the function and expression of MDR transporters in the adult BEC line, hCMEC/D3 and demonstrated an increase in P-gp activity following LPS (lipopolysaccharide, a bacterial infection mimic) treatment, with a concomitant decrease in *ABCB1* mRNA levels, and no change in protein expression. In the present study, protein levels were measured in whole cell lysate whereas functional assays target membrane P-gp activity. P-gp is known to be highly glycosylated ^42^ and a review by Czuba et al. ^57^ summarized different posttranslational modifications that regulate ABC transporters and outlined that phosphorylation of P-gp at serine residues S661, S667, S671, and S683 results in cell membrane trafficking of P-gp. Therefore, it is possible that posttranslational modifications are driving the increase in P-gp activity without changes in protein levels, and this will be addressed in future studies.

Functionally, an increase in P-gp activity in hfBECs in response to hypoxia, indicates greater fetal brain protection from xenobiotics and drugs that may be circulating in the fetal circulation, as well as alterations in the brain biodistribution of endogenous molecules like hormones and cytokines important for fetal brain development. Brain concentrations of the drug digoxin were found to be 35-fold higher in P-gp knockout mice compared to wild-type animals ^58^. Similarly, accumulation of other drugs including quinidine, cyclosporine, and erythromycin in the brain were observed in P-gp deficient animals, increasing brain toxicity ^59^. Alongside drug efflux, P-gp also regulates transport of endogenous factors that may be important for fetal brain development. The cytokine IL-6 is critical for fetal neurodevelopment and is also a substrate of P-gp. IL-6 is known to improve survival of neurons in culture, protect neurons from excitotoxic and ischemic insults, and regulate the development of astrocytes ^60^. Endogenous glucocorticoids like cortisol and corticosterone are also P-gp substrates and an important developmental trigger for the fetal brain. Low glucocorticoid levels are important for normal brain development as the fetal brain is highly sensitive to glucocorticoid excess ^32^. Early exposure to high glucocorticoid levels can affect growth as well as long-term endocrine and behavioral functions in offspring ^6^. As such, since fetal hypoxia induced by a reduction in maternal oxygen or by restriction of uteroplacental blood flow resulted in an increased in ACTH and cortisol concentrations in fetal plasma ^61^, therefore abundance of P-gp at the fetal BBB may help prevent glucocorticoids from entering and accumulating in the brain, thereby potentially maintaining adaptive brain homeostasis during hypoxia.

BCRP function was also increased by severe hypoxia in mid-gestation hfBECs, but only following extended exposure. BCRP transports a number of important substrates including endogenous molecules like folic acid. Folates are particularly important during rapid tissue growth and development, and hence it is of major importance during morphogenesis and organogenesis. Folate deficiency during pregnancy can lead to adverse outcomes such as neural tube defects and preterm birth^62^. Thus, specific endogenous BCRP substrates play an important role during brain development, and an increase in its activity following severe and chronic hypoxia indicates an increase in the efflux of such molecules out of the developing brain. Whether this adaptive response is protective or detrimental is unknown and requires further investigation.

### Limitations of the study

Potential limitations comprise the relatively low number of subjects (n=6/group) enrolled in the present study. This is due to the fact that availability of human fetal brain specimens for research is extremally limited. However, previous research investigating drug transporter expression and function in developing tissues has utilized a sample size of six per group in human ^8,47,48,50,63–65^ and animal ^66–69^ studies. As mandated by the REB, we are unable to retrieve clinical information on the fetal tissues collected. In this connection, maternal clinical information such as age, BMI, ethnicity, parity, and fetal sex could explain, at least in part, for some of the within group variability detected in some of the results. Finally, our model system does not allow investigation of potential changes in blood flow, which are known to influence regulation of transporter function. Future studies incorporating flow in *in vitro* culture systems will further strengthen interpretation of our results ^70,71^. Notwithstanding, the present study is novel and shows that hypoxia modifies the expression and function of the two primary drug transporters in hfBECs, in an age-dependent manner.

## Conclusions

In summary, we have identified specific changes in P-gp and BCRP expression or function after short term and prolonged severe hypoxia in early and mid-gestation hfBECs. The disconnect between levels of P-gp and BCRP (mRNA, protein, and function) are most likely attributed to hypoxia-induced changes in posttranslational modifications and shuttling of transporter proteins to the cell membrane. Longer-term exposure to hypoxia (48-hours) significantly decreased BCRP protein and function in early gestation hfBECs. This implies that the fetal brain may become more vulnerable to exposure to potentially harmful BCRP substrates present in the maternal circulation, during early development, but may increase brain deposition of other endogenous substrates such as folate. In mid-gestation, severe hypoxia led to rapid and sustained increases in P-gp and BCRP function. Changes in P-gp activity after just 6-hours suggests the high sensitivity of P-gp to hypoxia. The increase in activity of both transporters would potentially decrease transfer of substrates, including xenobiotics, drugs, and endogenous substrates across the fetal BBB.

## Data Availability

All data generated or analyzed during this study are included in this article.

## Ethics approval and consent to participate

Human fetal brain tissue was collected by the Women’s and Infants’ Health BioBank program at Sinai Health following written informed consent using a protocol approved by the Research Ethics Boards of Sinai Health and the University of Toronto (#18-0057-E).

## Acknowledgments

The authors thank the donors, RCWIH BioBank, the Lunenfeld -Tanenbaum Research Institute, and the Mount Sinai Hospital/UHN Department of Obstetrics and Gynaecology for the human specimens used in this study.

## Author Contributions

Conceptualization, H.M, G.I., E.B. and S.G.M.; Methodology, H.M, G.I., P.L., E.B. and S.G.M..; Investigation, H.M. and P.L.; Writing – Original Draft, H.M.; Writing –Review & Editing, E.B. and S.G.M.; Funding Acquisition, E.B. and S.G.M.; Resources, S.G.M.; Supervision, E.B. and S.G.M.

## Supplementary Information

Additional file 1: Supplementary Figure 1: Validation of hypoxia and cell viability in early and mid-gestation human primary fetal brain endothelial cells (hfBECs). Supplementary Figure 2: Inhibitor controls for P-gp and BCRP substrate specificity. Supplementary Figure 3: Effect of hypoxia on esterase activity in mid-gestation human primary fetal brain endothelial cells (hfBECs).

