## Supplementary Figures for "Hypoxia modulates P-glycoprotein (P-gp) and breast cancer resistance protein (BCRP) drug transporters in brain endothelial cells of the developing human blood-brain barrier"

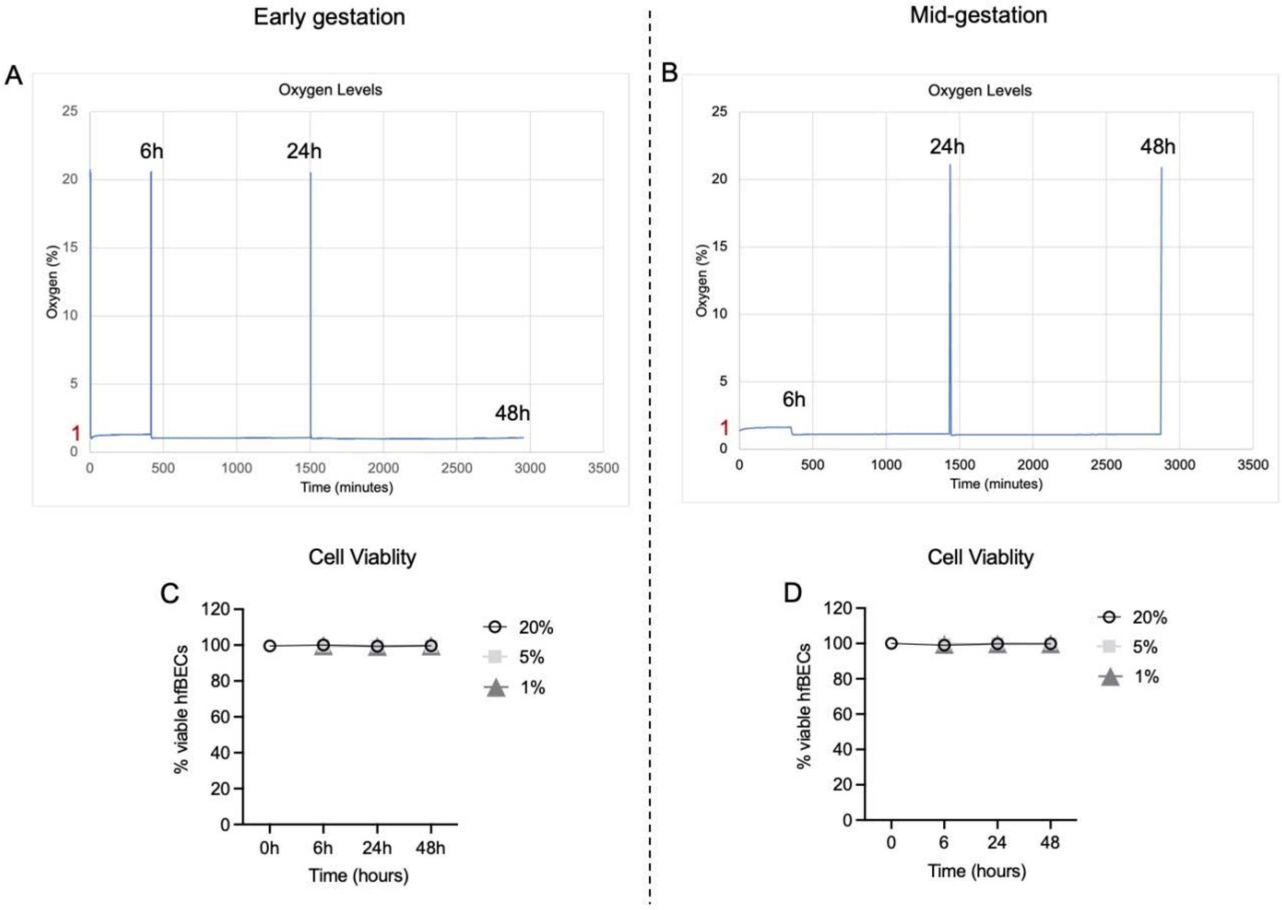


**Supplementary Figure 1: Validation of hypoxia and cell viability in early and mid-gestation human fetal brain endothelial cells (hfBECs)**. **(A-B)** Oxygen sensor measurements within the 1% oxygen chamber containing early and mid-gestation hfBECs. The peaks to 20% oxygen denote when the chamber was opened to take plates out for harvesting the cells after each respective time-point. **(C-D)** Cell viability was assessed by trypan blue dye exclusion test (*N*=6/group). Data are expressed as mean ± SEM.

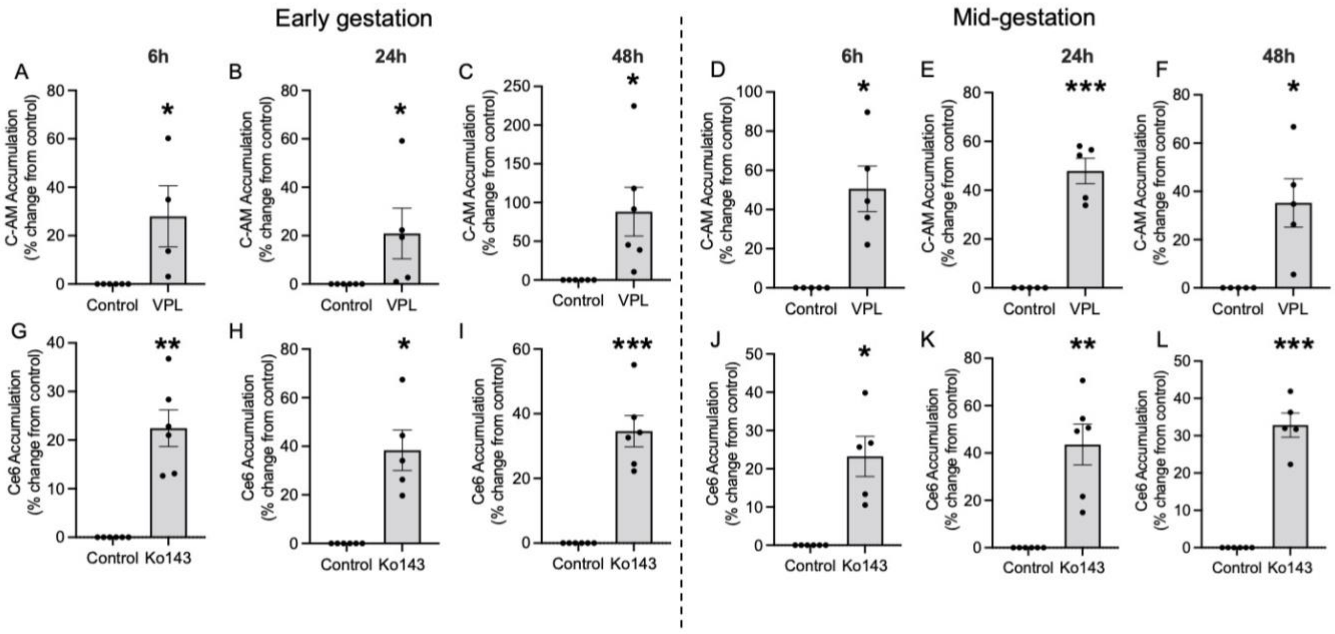


**Supplementary Figure 2: Inhibitor controls for P-gp and BCRP substrate specificity – early and mid-gestation human fetal brain endothelial cells (hfBECs).** The specificity of calcein-AM (C-AM) and chlorin e6 (Ce6) was determined using the P-gp inhibitor verapamil (VPL) and BCRP inhibitor Ko143, respectively. Accumulation of substrates was measured under 20% oxygen with and without the presence of inhibitor after **(A, D, G, J)** 6-hours, **(B, E, H, K)** 24-hours, and **(C, F, I, L)** 48-hours (*N*=4-6/group). Data are displayed as percent inhibition relative to control; increased accumulation of fluorescent substrates indicates reduced transporter activity. Statistical analysis: unpaired Student's t-test. Values are displayed as mean ± S.E.M. (*) $P<0.05$; (**) $P<0.01$; and (***) $P<0.001$.

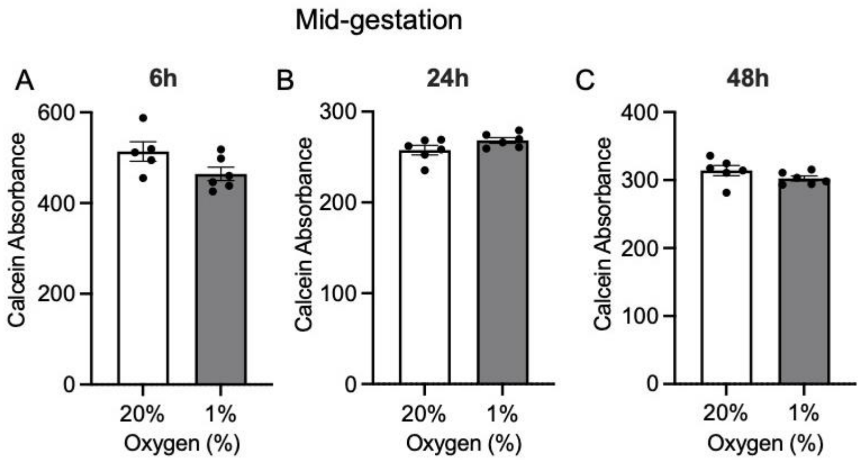


**Supplementary Figure 3: Hypoxia does not affect esterase activity in mid-gestation human fetal brain endothelial cells (hfBECs).** Relative fluorescence units (RFU) in lysed hfBECs following treatment with 1% oxygen compared to control after **(A)** 6-hours, **(B)** 24-hours, and **(C)** 48-hours (*N*=5-6/group). Statistical analysis: unpaired Student's t-test. Values are displayed as mean ± S.E.M.
